# The Impact of Brief Monocular Retinal Inactivation on the Central Visual System During Postnatal Development

**DOI:** 10.1101/2022.08.08.503194

**Authors:** Kevin R. Duffy, Nathan A. Crowder, Arnold J. Heynen, Mark F. Bear

**Affiliations:** Department of Psychology & Neuroscience, Dalhousie University, Halifax, NS, Canada, B3H 4R2; Picower Institute for Learning and Memory, Department of Brain and Cognitive Sciences, Massachusetts Institute of Technology, Cambridge, MA 02139

**Keywords:** monocular deprivation, monocular inactivation, dLGN, visual cortex, tetrodotoxin, neural plasticity, critical period

## Abstract

During a critical period of postnatal life, monocular deprivation (MD) of kittens by eyelid closure reduces the size of neurons in layers of the dorsal lateral geniculate nucleus (dLGN) connected to the deprived eye, and shifts cortical ocular dominance in favor of the non-deprived eye, modeling deprivation amblyopia in humans. Following long-term MD, temporary retinal inactivation of the non-deprived eye with microinjection of tetrodotoxin can promote superior recovery from MD, and at older ages, in comparison to conventional occlusion therapy. This suggests that monocular inactivation (MI) is a more potent approach to producing neural plasticity than occlusion. In the current study we assessed the modification of neuron size in the dLGN as a means of measuring the impact of a brief period of MI imposed at different ages during postnatal development. The biggest impact of inactivation was observed when it occurred at the peak of the critical period for ocular dominance plasticity. The effect of MI was evident in both the binocular and monocular segments of the dLGN, distinguishing it from MD that produces changes only within the binocular segment. With increasing age, the capacity for inactivation to alter postsynaptic cell size diminished but was still significant beyond the classic critical period. In comparison to MD, inactivation consistently produced effects that were about double in magnitude, and inactivation exhibited efficacy to produce neural modifications at older ages than MD. Notwithstanding the large neural alterations precipitated by inactivation, its anatomical effects were remediated with a short period of binocular visual experience, and vision through the previously inactivated eye fully recovered after washout of TTX. Our results demonstrate that MI is a potent means of modifying the visual pathway, and does so beyond the age at which occlusion is effective. The magnitude and longevity of inactivation to evoke neural modification highlights its potential to ameliorate disorders of the visual system such as amblyopia.

## Introduction

An important factor in the early postnatal development of visual neural circuits is provision of normal binocular vision. Obstruction of clear and balanced vision during a formative and highly plastic stage of development, called the critical period, can produce a rearrangement of neural connections after which the visual system exhibits abnormal responses to visual stimulation. Following even a brief period of MD by eyelid closure, physiological responses in the visual cortex skew in favor of the non-deprived eye, leaving the deprived eye able to control the activity of few cortical neurons (Wiesel and Hubel, 1963a; Hubel et al., 1977; Mioche and Singer, 1989). In cats and monkeys, the physiological shift in cortical ocular dominance elicited by MD is accompanied by anatomical modifications within binocular visual cortex that include a retraction of deprived-eye geniculostriate axon terminals (Shatz and Stryker, 1978; LeVay et al., 1980; Antonini and Stryker, 1993). This simplification of axons in visual cortex correlates with a reduction in the volume of their parent somas in the dLGN. Because cell body shrinkage is only observed in regions of the dLGN serving the binocular visual field (the “binocular segment” of the nucleus), it is not believed to be a consequence of altered retinal (orthodromic) activity, per se. Rather, reduced soma size after MD is considered to be a retrograde reflection of interactions between converging inputs from the two eyes in binocular visual cortex (Guillery and Stelzner, 1970; Guillery, 1972; Wiesel and Hubel, 1963b). This interpretation is supported by the finding that binocular competition in the control of dLGN cell size depends on synaptic plasticity within visual cortex (Bear and Colman, 1990).

The neural aberrations that arise from MD are thought to represent the core pathology of an accompanying visual impairment - deprivation amblyopia, that is characterized by a pronounced reduction in spatial acuity (Giffin and Mitchell, 1978) as well as loss of binocularity and stereopsis (Timney, 1983; Vorobyov et al., 2007). Reversal of the effects of MD is possible during the highly plastic critical period if the deprived eye is opened and the originally nondeprived eye is closed (Blakemore and Van Sluyters, 1974; Movshon, 1976), a procedure referred to as reverse occlusion that is analogous to human full-time patching for the treatment of amblyopia. As with patching, reverse occlusion applied at ages beyond the critical period when plasticity capacity is low has limited potential to elicit recovery from the structural and functional consequences of a prior period of MD (Wiesel and Hubel, 1965; Blakemore and Van Sluyters, 1974). This age-related recalcitrance to recovery is also observed in humans, and is of particular concern for amblyopia produced by deprivation because of a more restricted window for recovery in comparison to other forms of the disorder (Birch and Stager, 1998; Holmes and Levi, 2018).

Recent mouse and cat studies have demonstrated that post-critical period recovery from MD can occur if the dominant eye’s retina is temporarily inactivated with intravitreal microinjection of tetrodotoxin (TTX), a voltage-gated sodium channel blocker that can reversibly eliminate retinal ganglion cell activity (Duffy et al., 2018; Fong et al., 2021). Following a period of MD, inactivation of the dominant (fellow) eye restores neural responses to the weaker one, and promotes recovery from the MD-induced atrophy of neuron soma size in the dLGN. Importantly, this recovery from the effects of MD occurs at an age when reverse occlusion is incapable of promoting significant recovery from the original deprivation event (Blakemore and Van Sluyters, 1974; Duffy et al., 2018). While the mechanisms that elicit recovery following inactivation remain unknown, and are currently being explored, the superior efficacy of inactivation as a recovery strategy relative to occlusion therapy suggests that the two procedures are not equal in their impact on the visual system.

An objective of the current study was to investigate potential differences between these two therapeutic approaches (temporary form deprivation versus inactivation). We chose to measure an anatomical hallmark of visual deprivation as a means of probing the impact of both procedures and assessing potential differences between monocular lid closure and monocular retinal inactivation, with an ultimate goal to provide insight into the superior efficacy of inactivation to promote neural recovery. One possibility is that MI produces effects in the visual system that mirror those of MD but are simply greater in magnitude. An alternative possibility is that neural modifications elicited by the two forms of visual deprivation exhibit similarities but are on balance fundamentally different. This latter possibility is consistent with an earlier study that demonstrated the two conditions are different in their impact on visual cortex plasticity (Rittenhouse et al., 1999).

For over 50 years, measurement of neuron soma size within the cat dLGN has provided a reliable, robust, and sensitive means of assessing the impact of MD on the structure and function of neurons in the primary visual pathway (Wiesel and Hubel, 1963b; Guillery and Stelzner, 1970; Dursteler and Movshon, 1976; Kutcher and Duffy, 2007). In this study, we compared the effects of MD to those of retinal inactivation by measuring the magnitude and retinotopic location of modifications within the dLGN across different ages in development. The eye-specific lamination and retinotopic organization of the cat dLGN make it ideal for selective quantification of both magnitude and topography of neural changes elicited by the two forms of monocular manipulation. Importantly, this experimental design enabled within-animal comparisons between affected and normal-eye layers that can subvert complications arising from issues such as high variability between animals (Guillery, 1973), and which is particularly useful in studies using higher mammals where low subject numbers are typical.

## Methods

### Animals

Anatomical studies were conducted on 15 cats that were all born and raised in a closed breeding colony at Dalhousie University. Rearing and experimental procedures were conducted in accordance with protocols approved by the University Committee on Laboratory Animals at Dalhousie, and conformed to guidelines from the Canadian Council on Animal Care. Tissues from some of the animals in this investigation were collected for previous studies, and samples from these animals were obtained from our cat brain tissue bank. We examined dLGN slices from three groups of animals whose rearing histories are detailed in Table 1. Animals were reared with MD by eyelid closure (n=5; 3 males, 2 females), or with MI (n=8; 4 males, 4 females), or received MI followed by a period of binocular vision (n=2; 1 male, 1 female).

Tissues from animals in our MD groups were obtained from our brain bank and had the eyelids of one eye closed for 14 days starting either at postnatal day (P) 30, the peak for ocular dominance plasticity, or at later ages (P42 and P70) when the efficacy of MD to shift ocular dominance has diminished (Olson and Freeman, 1980). This 14-day MD duration was selected as it represented the closest available deprivation duration relative to our retinal inactivation group. The comparison group of animals had one eye inactivated for 10 days starting either at P30, P42, P70, or 22 weeks of age.

### Monocular Deprivation

Animals were monocularly deprived under general gaseous anesthesia (3-4% isoflurane in oxygen) and the procedure involved closure of the upper and lower palpebral conjunctivae of the left eye with sterile 5-0 vicryl, followed by closure of the eyelids with 5-0 silk suture. Upon completion of the procedure, animals were administered Metacam (0.05 mg / kg) for postprocedure analgesia, local anesthesia was produced with application of Alcaine sterile ophthalmic solution (1% proparacaine hydrochloride; CDMV, Canada), and a broad-spectrum topical antibiotic (1% Chloromycetin; CDMV) was administered to mitigate infection after surgery. Quality of the eye closure was monitored daily to ensure the lids were in good health and fully closed.

### Retinal Inactivation

Retinal inactivation was performed under general anesthesia with 3-4% isoflurane in oxygen. Consistent with our previous studies, animals had their right eye inactivated with intravitreal microinjection of TTX (ab120055; abcam, USA) that was solubilized in citrate buffer at 2mM. For each animal, dosage was scaled according to eye size (Thorn, 1976). We administered 0.5 μl of TTX per mm of vitreous chamber length. This approximate dosage blocks action potentials of affected cells without obstructing critical cellular functions such as fast axoplasmic transport (Ochs and Hollingsworth, 1971). Intravitreal injections were administered through a puncture made with a 30-gauge disposable sterile needle that produced a small hole in the sclera located at the pars plana. The measured volume of TTX solution was slowly dispensed into the vitreous chamber with a sterilized Hamilton syringe (Hamilton Company, USA) that had a fixed 30-gauge needle (point style 4). The needle was positioned through the original puncture and placed 5-10 mm into the chamber angled away from the lens. The total volume of TTX was dispensed slowly, then the needle was held in place for about a minute before it was retracted.

Following intraocular injection, topical antibiotic (T-1%; Aventix, Canada) and anesthetic (Alcaine; Alcon, Canada) were applied to the eye to prevent post-injection complications. Metacam (0.05 mg / kg) was administered for post-procedure analgesia. To achieve 10 days of retinal inactivation, animals received 5 injections, one every 48 hours, and for each injection the original puncture site was used to avoid having to make another hole. During the period of inactivation we confirmed inactivation by noting anisocoria due to pupil dilation in the inactivated eye, absence of a pupillary light reflex, and the lack of visuomotor behaviors such as visual placing, visual startle, and the ability to track a moving laser spot using the inactivated eye. These assessments were made while vision in the non-injected eye was briefly occluded with an opaque contact lens.

### Sample Preparation for Histology

In preparation for histology, animals were euthanized with a lethal dose of sodium pentobarbital (Pentobarbital Sodium; 150 mg/kg) and shortly after were exsanguinated by transcardial perfusion with approximately 150 ml of phosphate buffered saline (PBS) followed by an equal volume of PBS containing 4% dissolved paraformaldehyde. Brain tissue was immediately extracted and the thalamus was dissected from the remainder of the brain in order to prepare the dLGN for sectioning and histological processing. Tissue containing the dLGN was cryoprotected, then was cut coronally into 25-μm thick sections using a sliding microtome. Some of the sections obtained from our brain bank were cut at 50 μm. Tissue slices were stored at −20 degrees Celsius within an antigen preservative solution (Burke et al., 2009) until used for the study. Tissues examined in this study were all subjected to the same extraction and preparation procedures, the same storage settings, and sections from all animals were stained following the same protocol.

### Nissl Staining

For each animal, 6 sections containing the left and right LGN were mounted onto glass slides and stained with a 1% Nissl solution (ab246817; Abcam, USA). Stained sections were differentiated in 70% ethanol, then were dehydrated in a graded series of ethanols before clearing with Histo-Clear. Sections were then coverslipped with mounting medium (Permount; Fisher Scientific, Canada) and allowed to dry before microscopic evaluation.

### Anatomical Quantification

Measurements in this study were performed blind to each animal’s rearing condition. The cross-sectional area of neuron somata within A and A1 layers of the left and right dLGN was measured from Nissl-stained sections using the nucleator probe available on a computerized stereology system (newCAST; VisioPharm, Denmark). All area measurements were performed using a BX-51 compound microscope with a 60X oil-immersion objective (Olympus; Canada). Neurons were distinguished from glial cells using established selection criteria (Wiesel and Hubel, 1963a; Guillery and Stelzner, 1970) that included measurement of cells with dark cytoplasmic and nucleolar staining, and with light nuclear staining (Duffy et al., 2012). Adherence to these criteria permitted avoidance of cell caps and ensured that measurements were taken from neurons cut through the approximate somal midline. Approximately 500-1000 neurons were measured from each animal.

The impact of MD and MI was calculated using an ocular dominance index (ODI) that we have used previously (Duffy et al., 2014), and that revealed the percentage difference between eye-specific dLGN layers:

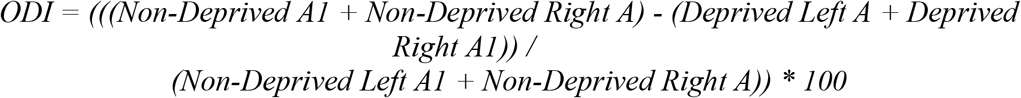

Statistical comparisons between eye-specific layers in each condition were made by treating the mean soma size of each layer as a single observation, then employing an unpaired (one-tailed) t-test to determine if affected neurons were smaller in size from those serving the normal eye. An exponential growth curve was used to characterize the effect of MI across age, and a Pearson product-moment correlation coefficient was used to calculate the goodness of fit.

### Physiology: Visually-Evoked Potentials

All cat recordings were conducted in anesthetized animals with full-field visual stimuli presented on an LCD monitor in the binocular visual field at a viewing distance of 70 cm. In preparation for each recording session, animals were anesthetized with 1-1.5% isoflurane, and supplemental sedation was provided with intramuscular acepromazine (0.06-0.1mg/kg). Hair on the head was trimmed and a disposable razor was used to shave parts of the scalp where recording sites were located, two positioned approximately 2-8 mm posterior and 1-4 mm lateral to interaural zero over the presumptive location of the left and right primary visual cortices, and another site over the midline of the frontal lobes that acted as a reference. Electrode sites were abraded with Nuprep EEG skin preparation gel (bio-medical, MI, USA), and were then cleaned with alcohol pads. Reusable 10 mm gold cup Grass electrodes (FS-E5GH-48; bio-medical) were secured to each electrode site using Ten20 EEG conductive paste (bio-medical, USA) that was applied to the scalp. Impedance of the recording electrodes was measured in relation to the reference electrode to ensure values for each were below 5 kΩ. Electrophysiological signals were amplified and digitized with an Intan headstage (RHD2132; 20kHz sampling frequency), then recorded using an Open Ephys acquisition board and GUI software (Open Ephys, USA)(Siegle et al., 2017). Stimuli were generated using custom software developed in Matlab using the Psychophysics Toolbox extension (Brainard, 1997; Pelli, 1997), and consisted of full contrast square wave gratings with a 2 Hz contrast reversal frequency (Bonds, 1984; Norcia et al., 2015; Pang and Bonds, 1991). Blocks of grating stimuli at different spatial frequencies (0.05, 0.1, 0.5, 1, and 2 cycles per degree (cpd)) or a luminance-matched grey screen were presented in pseudo-random order for 20 s each, with the grey screen also displayed during a 2 s interstimulus interval. Each block was repeated at least 6 times. Each eye was tested in isolation by placing a black occluder in front of the other eye during recording. Eyes were kept open with small specula, and the eyes were frequently lubricated with hydrating drops. Recording sessions lasted about 1 hour and animal behavior was observed for at least an additional hour post-recording to ensure complete recovery.

### Physiological Quantification

The raw electroencephalogram was imported to MATLAB where it was high-pass filtered above 1 Hz, then subjected to Fourier analysis (Bach and Meigen, 1999; Norcia et al., 2015). The magnitude of VEPs was calculated as the sum of power at the stimulus fundamental frequency plus 6 additional harmonics (2-14 Hz; DiCostanzo et al., 2020). Baseline nonvisual activity was calculated as the sum of power at frequencies 0.2 Hz offset from the visual response (2.2-14.2 Hz). To assess possible differences in VEP power for each eye across experimental conditions, a repeated measures ANOVA was employed using Holm-Sidak’s multiple comparisons test treating measurements from the right and left V1 across 0.05 cpd and 0.1 cpd stimuli as single observations.

## Results

### Monocular Deprivation

We started our investigation by measuring the effect of a 14-day MD within the dLGN. This duration of MD was selected as a comparison to the effect of 10 days of MI because among our brain bank samples this period of MD, while slightly longer, was the closest match. Imposition of a 14-day MD starting at the peak of the critical period (P30) produced a marked reduction in the size of neurons within deprived dLGN layers that was easily appreciated upon microscopic examination (**Figure 1A**). Consistent with findings from Guillery and Stelzner (1970), deprived neurons appeared obviously smaller than non-deprived counterparts only within the binocular segment of the dLGN layers (**Figure 1B**), where axons of neurons serving the left and right eye are subjected to competition at the level of V1 (Guillery, 1972). Within the laterally-positioned segment of the A layers, where neurons project to the monocular crescent representation of V1, there was no apparent difference between non-deprived and deprived soma size (**Figure 1C**). Stereological quantification of neuron cross-sectional soma area measured from the binocular segment revealed that deprived neurons (mean = 190μm^2^; SD = 9μm^2^) were 25% smaller than non-deprived neurons (mean = 253μm^2^; SD = 14μm^2^; **Figure 1D**), and this difference was significant (*t* = 7.498, d.f. = 6, p < 0.001). However, within the monocular segment, deprived (mean = 195μm^2^; SD =13 μm^2^) and non-deprived (mean = 185μm^2^; SD = 12μm^2^) neurons were different by only 5% (**Figure 1E**) and this was not a statistically significant difference (*t* = 0.847, d.f. = 2, p > 0.05).

**Figure 1.**
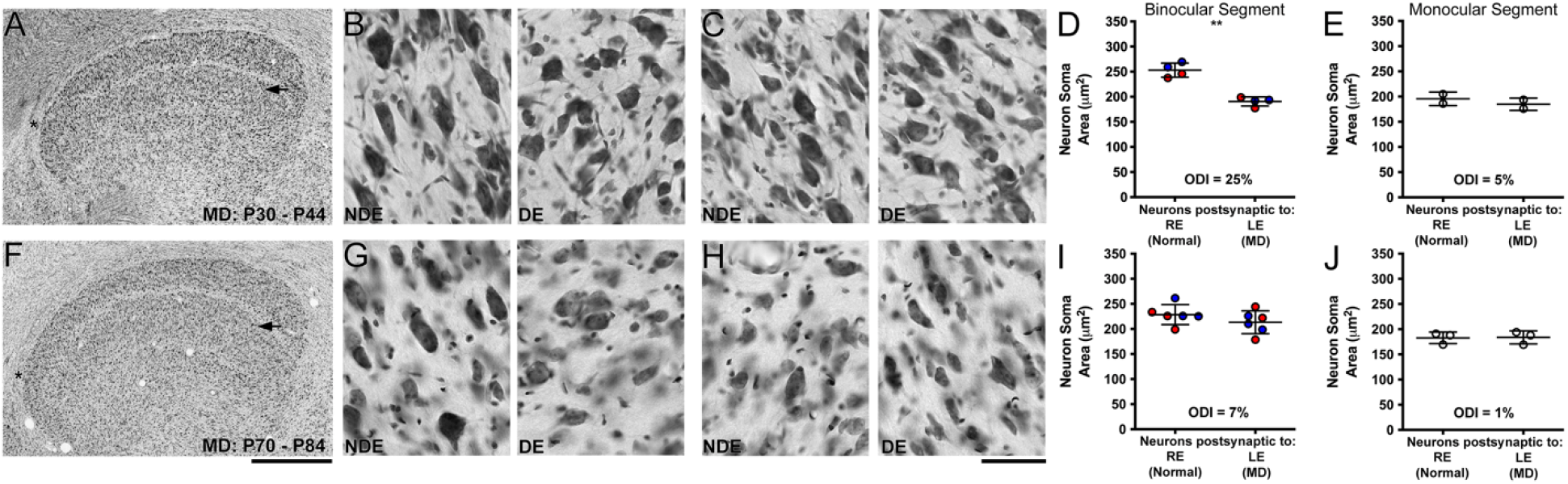
The impact of a 14-day MD imposed at or beyond the critical period peak. Low magnification view of the left dLGN stained for Nissl substance from an animal that received 14 days of MD at 4 weeks of age (A) revealed a reduction of neuron soma size within the deprived layer (arrow). At higher magnification the difference between non-deprived eye (NDE) and deprived eye (DE) neuron size was evident in the binocular segment of the dLGN (B) but not within the monocular segment (C). Quantification of neuron soma area revealed a 25% reduction in the size of deprived neurons within the binocular segment (D), but this effect did not extend into the monocular segment where eye-specific neurons were comparable in size (E). A much smaller effect on dLGN cell size was observed in the deprived layer (arrow) when 14 days of MD was imposed at 10 weeks of age (F), and this was reflected by observation at higher magnification with no obvious alteration in neuron soma size within either the binocular (I) or monocular (J) segments. These observations were supported with quantification that revealed a diminished impact on neuron size when MD occurred at 10 weeks of age. Scale bars = 1 mm (A and F) and 50 μm (B, C and G, H). Red and blue data points indicate measurements from A and A1 layers, respectively. Asterisks in A and F indicate location of the monocular segment in the A layer. The timing of MD is shown in the lower left of A and F. Double asterisks indicate statistical significance below 0.05.

When the same 14-day MD was imposed later in development, at 10 weeks of age, we observed a smaller effect in the dLGN (**Figure 1F**). At this older age, deprived neurons within the binocular segment were only slightly smaller than non-deprived neurons (**Figure 1G**), and no difference in size was noted for neurons located within the monocular segment (**Figure 1H**). Quantification of neuron soma size reflected our observation that at 10 weeks of age, MD for 14 days had a reduced effect within the binocular segment (deprived: mean = 213μm^2^, SD = 23μm^2^; non-deprived: mean = 228μm^2^; SD = 20μm^2^; **Figure 1I**), and virtually no effect within the monocular segment (deprived: mean = 184μm^2^, SD = 13μm^2^; non-deprived: mean = 183μm^2^; SD = 12μm^2^; **Figure 1J**). The difference between deprived and non-deprived neuron size within the binocular segment was 7% but not significantly different (*t* = 1.236, d.f. = 10, p > 0.05), and deprived and non-deprived neurons within the monocular segment were different by only 1% and also not significantly different (*t* = 0.096, d.f. = 4, p > 0.05). These results demonstrated that at this older age, MD had a diminished and insignificant impact on the size of dLGN neurons.

### Retinal Inactivation

The effect of monocular retinal inactivation was quantitatively and qualitatively different than that produced by MD. Monocular retinal inactivation for 10 days that began at the critical period peak produced an obvious alteration within dLGN layers connected to the inactivated eye, and this was clearly visible at low magnification in the two animals we examined (**Figure 2A** and **D**). Within the binocular segment, inactivated layers contained Nissl-stained neurons that were 44% smaller (mean = 156μm^2^, SD = 16μm^2^) and expressed a pale appearance relative to neurons located within normal-eye layers (mean = 278μm^2^, SD = 22μm^2^; **Figure 2B** and **C**). Contrary to what was observed in the monocular segment after MD, neurons within the monocular segment serving the inactivated eye were noticeably smaller (mean = 184μm^2^, SD = 24μm^2^) and paler in comparison to counterparts from the normal-eye layers (mean = 229μm^2^, SD = 20μm^2^; **Figure 2E**). Whereas after MD there was no soma size difference between deprived and non-deprived neurons within the monocular segment, following 10 days of MI we observed that inactivated-eye neurons were distinctly smaller than normal-eye neurons, which was quantified as a 20% difference (**Figure 2F**). The observed reduction of soma size after inactivation was statistically significant for both the binocular (*t* = 11, d.f. = 10, p < 0.001) and monocular (*t* = 2.5, d.f. = 4, p < 0.05) segments.

**Figure 2.**
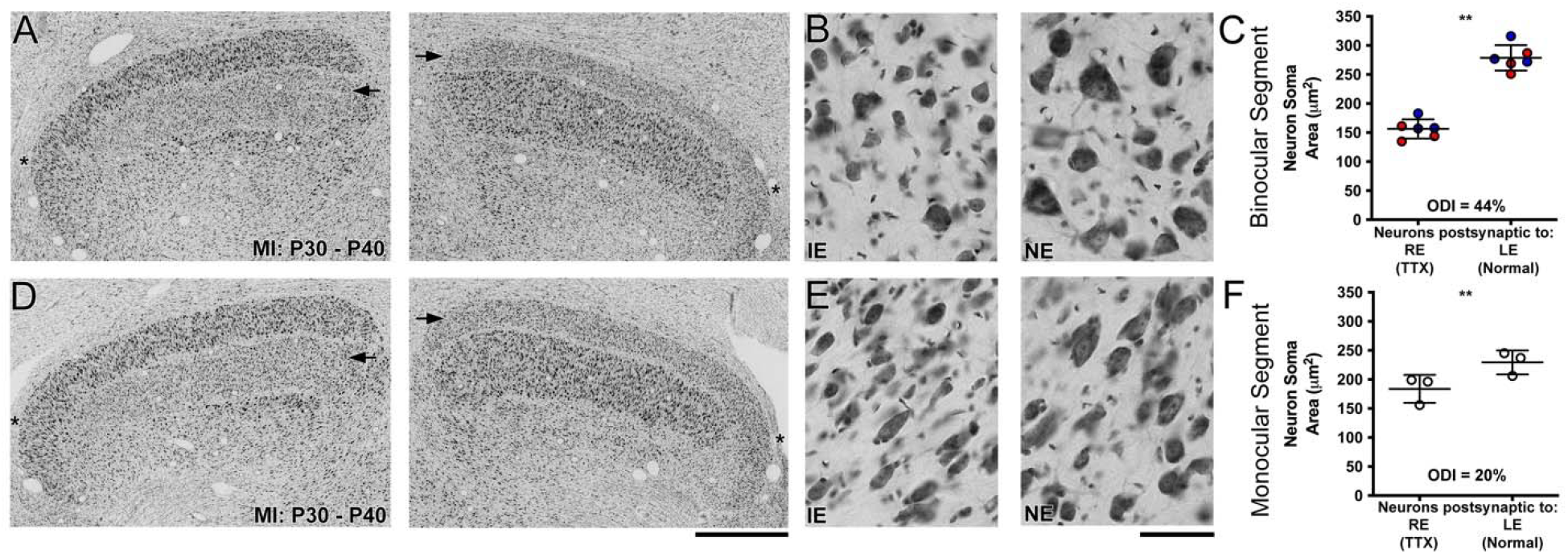
The effect of 10 days of retinal inactivation imposed at 4 weeks of age, the peak of critical period in cats (Olson and Freeman, 1980). Microscopic inspection of the left and right dLGN from two animals revealed a substantial loss of Nissl staining within layers serving the inactivated eye (IE; arrows in A and D) relative to those of the normal eye (NE), and this effect was accompanied by a sizable reduction of neuron soma area within both the binocular (B) and monocular (E) segments. Stereological quantification of neuron soma area confirmed these observations by showing that inactivated-eye neurons were 44% smaller than those serving the normal eye within the binocular segment (C), which was about double the effect size in comparison to MD. Distinct from the effects of MD, retinal inactivation rendered inactivated neurons 20% smaller within the dLGN’s monocular segment. Scale bars = 1 mm (A and D) and 50 μm (B and E). Red and blue data points indicate measurements from A and A1 layers, respectively. Asterisks in A and D indicate location of the monocular segment in the A layers. The timing of MI is shown in the lower left of A and D. Double asterisks indicate statistical significance below 0.05.

As we increased the age at which retinal inactivation began, the impact on neuron size within the dLGN was reduced. Although ten days of inactivation started at 10 weeks of age had a noticeable effect on soma size (**Figure 3A** and **D**), this was diminished in comparison to what was observed when the same period of inactivation occurred at 4 weeks of age. At 10 weeks of age, neurons within the binocular segment connected to the inactivated eye were clearly smaller but did not exhibit the obvious staining pallor that was observed after inactivation at the critical period peak (**Figure 3B**). Within the monocular segment (**Figure 3E**), a reduction of soma size was likewise evident within inactivated-eye layers; however, the effect magnitude did not appear to be as reduced by age. In other words, unlike the binocular segment, inactivated-eye neurons within the monocular segment showed only a slight reduction in effect magnitude in comparison to that measured following inactivation at the critical period peak. Quantification of soma area supported these observations by showing that inactivated-eye neurons in the binocular segment were rendered 23% smaller (inactivated: mean = 207μm^2^, SD = 3μm^2^; normal: mean = 272μm^2^; SD = 18μm^2^; **Figure 3C**), which was about matched in magnitude by the monocular segment that showed an 18% difference in soma size (inactivated: mean = 202μm^2^, SD = 5μm^2^; normal: mean = 249μm^2^; **Figure 3F**). The measured reduction of neuron soma size was significant for both the binocular (*t* = 7.242, d.f. = 10, p < 0.001) and monocular (*t* = 6.453, d.f. = 2, p < 0.05) segments. These results indicated that with increasing age the effect of retinal inactivation was reduced in magnitude, but the effect was still considerably larger than that elicited by a similar period of MD imposed at the same age. That the impact of inactivation was similar between the binocular and monocular segments suggests that the effect at this age was not driven by competition between geniculocortical afferents serving the two eyes.

**Figure 3.**
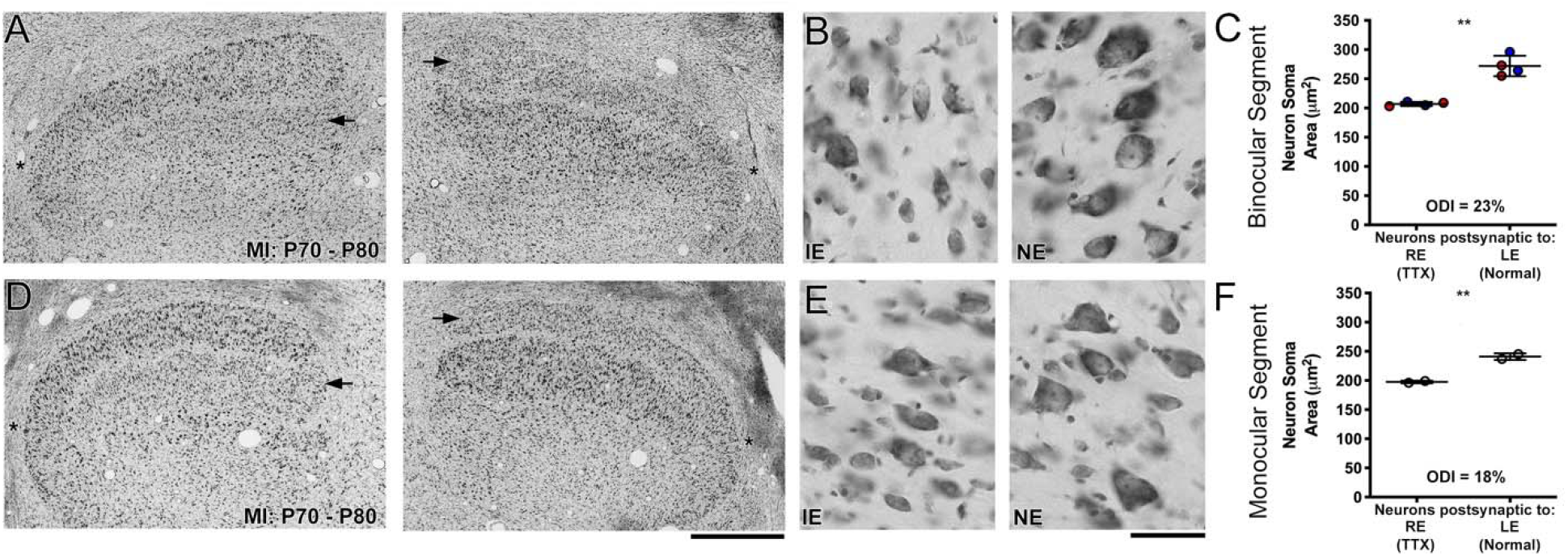
The impact of 10 days of retinal inactivation started at 10 weeks of age when MD has a negligible effect on dLGN cell size. Examples of Nissl staining in the right and left dLGN from two animals revealed a noticeable difference in layers serving the inactivated eye (arrows). This was better appreciated at higher magnification, which showed that neurons were smaller within inactivated-eye layers compared to those of the fellow eye. This difference in neuron size was apparent in both the binocular (B) and monocular segments (E). Quantification of neuron soma area supported these observations by showing that inactivated-eye neurons within the binocular segment were 23% smaller than those serving the normal eye (C), and 18% smaller within the monocular segment. Scale bars = 1 mm (A and D) and 50 μm (B and E). Red and blue data points indicate measurements from A and A1 layers, respectively. Asterisks in A and D indicate location of the monocular segment in the A layers. The timing of MI is shown in the lower left of A and D. Double asterisks indicate statistical significance below 0.05.

The oldest animals in this study had one eye inactivated for 10 days starting at 22 weeks of age (**Figure 4**). Ten days of MI at this age produced an effect on soma size that was not easily appreciated at low magnification (**Figure 4A** and **D**). Even at high magnification, the effect of inactivation was not as obvious as it was at the younger ages we examined, and this was true for both the binocular and monocular segments that were not appreciably different (**Figures 4B** and **E**). Stereological quantification of soma size matched our qualitative observations by showing a small difference between eye-specific neurons within the binocular (**Figure 4C**) and monocular (**Figure 4F**) segments. Neurons within inactivated-eye layers of the binocular segment (mean = 241μm^2^, SD = 13μm^2^) were 10% smaller than neurons in normal-eye layers (mean = 269μm^2^, SD = 17μm^2^), and this was a statistically significant difference (*t* = 2.59, d.f. = 6, p < 0.05). A similar result was obtained from the monocular segment where neurons were different by 8% (inactivated: mean = 229μm^2^, SD = 10μm^2^; normal: mean = 250μm^2^, SD = 9 μm^2^; **Figure 4F**); however, this was not significantly different (*t* = 2.237, d.f. = 2, p > 0.05).

**Figure 4.**
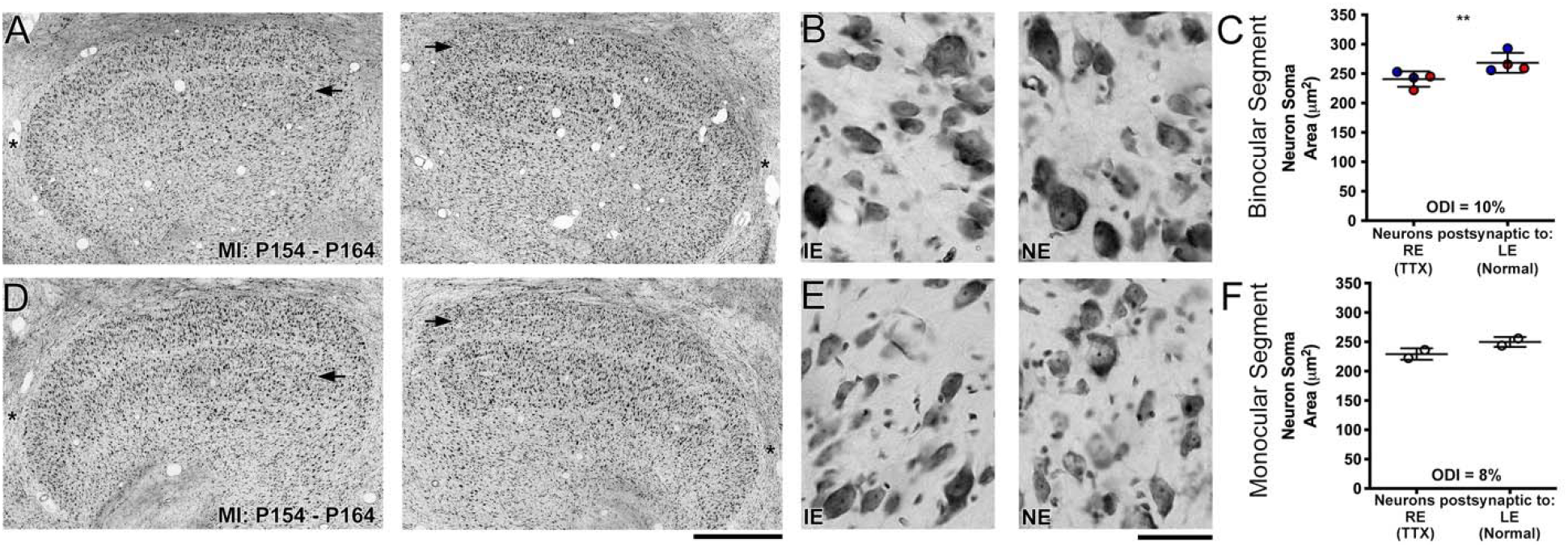
The consequence of 10 days of retinal inactivation started at 22 weeks of age when the classical critical period has passed. Low magnification examples of Nissl staining within the right and left dLGN from two animals (A and D) revealed a minor change within the layers serving the inactivated eye (arrows). At higher magnification it was also obvious that neurons were only slightly smaller within inactivated-eye layers compared to those serving the felloweye, which was evident within both the binocular (B) and monocular (E) segments. Quantification of neuron soma area mirrored these observations by showing that inactivated-eye neurons in the binocular segment were only 10% smaller than those serving the normal eye (C), and within the monocular segment the difference was similar at 8%. Scale bars = 1 mm (A and D) and 50 μm (B and E). Red and blue data points indicate measurements from A and A1 layers, respectively. Asterisks in A and D indicate location of the monocular segment in the A layers. The timing of MI is shown in the lower left of A and D. Double asterisks indicate statistical significance below 0.05.

To illustrate the impact of imposing 10 days of MI at different postnatal ages, we plotted the percentage difference in soma size between inactivated and normal-eye dLGN layers for the binocular and monocular segments across the ages examined (**Figure 5**). Within the binocular segment, there was a progressive decline in the effect of retinal inactivation with increasing age (**Figure 5A**). From the peak of the critical period to 22 weeks of age, the effect of inactivation was reduced by about 75%, from an average effect size of 44% down to 10%. Notably, even at 22 weeks of age, the oldest age we examined, a small residual effect of inactivation was observed. The decline in susceptibility to inactivation with age was well characterized by an exponential growth curve (R^2^ = 0.97). The effect of MI was consistently about double the magnitude of that produced by MD within the binocular segment. The effect of inactivation within the monocular segment was also reduced with age (**Figure 5B**), and the data were also fit with an exponential growth curve (R^2^ = 0.72). Diminution of the effect of inactivation with increasing age was less in the monocular segment compared to the binocular segment, which was based on the observation that effect magnitude in the monocular segment at 4 and 10 weeks of age was similar. In aggregate, results from our MI groups highlight distinct differences from MD that include an abiding efficacy to elicit cell size changes at older ages, as well as an ability to produce alterations within the monocular segment of the dLGN.

**Figure 5.**
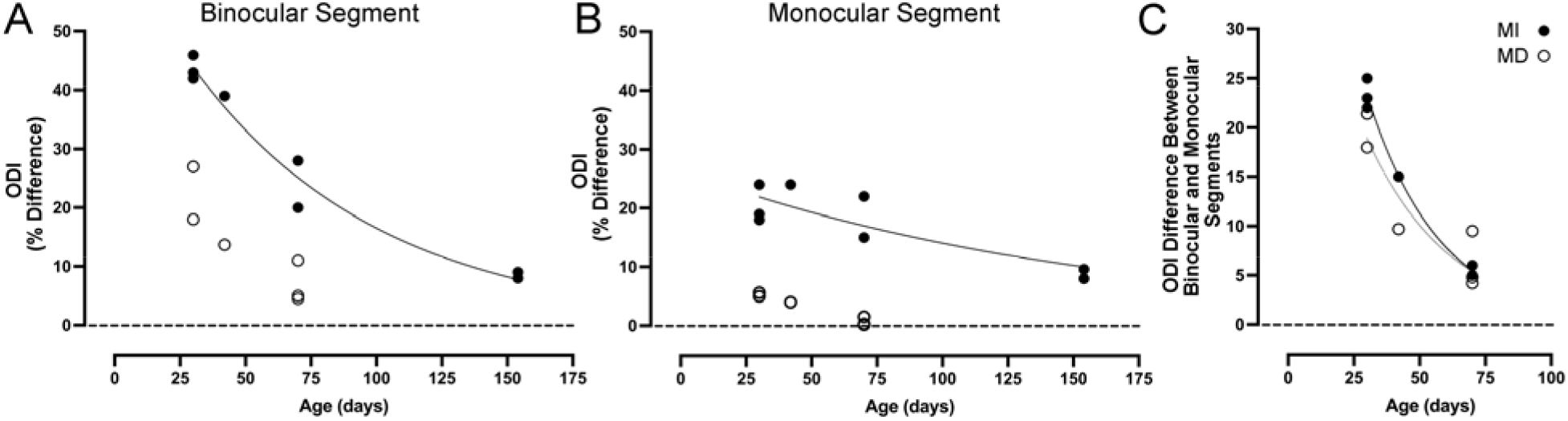
The effect of a fixed duration of MI imposed at different ages across postnatal development. Within the binocular segment, the effect of 10 days of inactivation was clearly reduced with age (A). Between 4 and 22 weeks of age, there was a 75% reduction in the efficacy of inactivation, though, importantly, a small effect still remained even at the oldest age examined. In comparison to MD, MI elicited greater change within the dLGN, yielding effects that were about double in magnitude. Within the monocular segment (B), the impact of inactivation was likewise reduced with age but less so in comparison to the binocular segment. A clear distinction between the effect of MD and inactivation was observed within the monocular segment: whereas the size of neurons in the monocular segment exhibit little change with MD (Guillery and Stelzner, 1970; Hickey et al., 1977), inactivation had an effect on neuron size even at oldest age examined. Subtraction of the differences in ODI within the monocular segment from the differences in the binocular segment revealed that the developmental profiles for MD and MI groups were similar (C), suggesting the effect of MD and MI mediated by binocular competition is comparable.

Whereas MD applied at 10 weeks of age produced only a small change in the dLGN, retinal inactivation applied at this age reduced soma size significantly, and by an amount that was comparable across the binocular and monocular segments. Uniformity of the response across segments raises the possibility that the effect of retinal inactivation at this age is driven by something other than binocular competition. Indeed, when we subtracted the observed differences in ODI within the monocular segment (possibly related to diminished trophic support from the retinogeniculate afferents) from the differences in the binocular segment (that include the additive retrograde effect of binocular competition in visual cortex), the developmental profiles for MD and MI groups were similar (**Figure 5C**). This comparison is thought to isolate the effect of binocular competition (Sherman and Spear, 1982), and therefore showed that MI and MD are comparable in this regard. On the other hand, the interruption of supposed retinogeniculate influence is clearly different, as evidenced by the effect observed in the monocular segment. Just as an inactive muscle fiber rapidly recovers size when the temporarily silenced motoneuron activity is restored, we wondered if dLGN cell size changes ascribed to reduced retinal activity is similarly transient. In previous studies, we demonstrated that fellow-eye retinal inactivation applied at 10 weeks of age can promote recovery from the effects of a prior long-term MD, and critically the inactivation produced no lasting detriment to the inactivated eye or to the dLGN layers connected to it (Duffy et al., 2018; Fong et al., 2021; DiCostanzo et al., 2020). We were therefore interested to test the durability of the effect of retinal inactivation by providing a subset of animals with binocular vision after they received 10 days of retinal inactivation applied at 10 weeks of age. After allowing the period of inactivation to wear off, layers of the dLGN connected to the previously inactivated eye had a normal appearance with no evidence of staining pallor or neuron atrophy within inactivated-eye layers (**Figure 6A and D**). These observations were confirmed with high magnification images that showed comparable staining characteristics between eye-specific layers within the binocular (**Figure 6B**) and monocular (**Figure 6E**) segments of the dLGN. Quantification of soma size indicated that the change in neuron size provoked by unilateral retinal inactivation (**Figure 3**) was erased with provision of 10 days of binocular vision. Within the binocular segment, there was only a 5% difference in the size of neurons (inactivated: mean = 229μm^2^, SD = 10μm^2^; normal: mean = 250μm^2^, SD = 9; **Figure 6C**), and this was not a significant difference (*t* = 0.674, d.f. = 6, p > 0.05). Similarly, only a 4% difference in neuron soma size was observed within the monocular segment, which also was not significantly different (inactivated: mean = 229μm^2^, SD = 10μm^2^; normal: mean = 250μm^2^, SD = 9; *t* = 0.068, d.f. = 2, p > 0.05; **Figure 6F**). These results indicate that the effect of MI on dLGN soma size is ephemeral, at least when it is administered at an age beyond the critical period peak.

**Figure 6.**
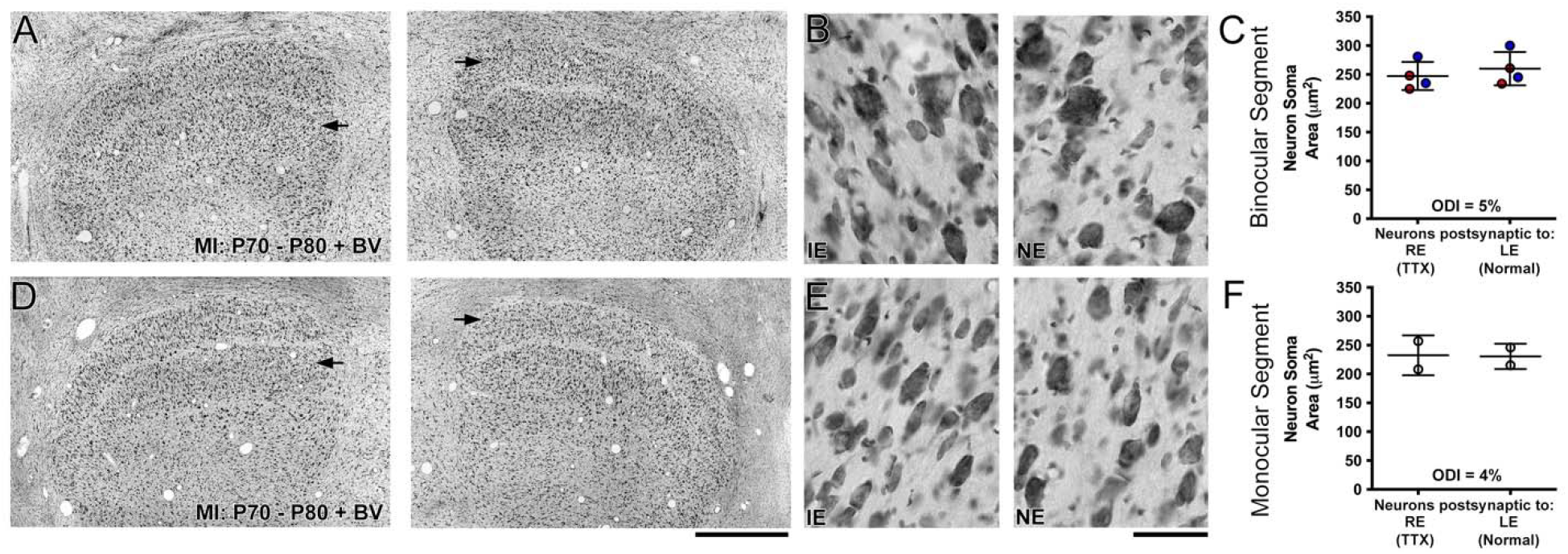
Recovery from the effects of 10 days of retinal inactivation applied at 10 weeks of age. Eye-specific layers within the binocular segment of the dLGN were not obviously different in staining characteristics when MI was immediately followed by 10 days of normal binocular vision (A and D). Previously inactivated layers (arrows) contained neurons that were comparable in size to those within layers serving the normal eye, and this was also evident at high magnification for both the binocular (B) and monocular (E) segments. Quantification of neuron soma area revealed balance between the eye-specific layers that was comparable to normal (Fong et al., 2016), and this was true for measurements from the binocular (C) and monocular (F) segments. This indicates that the modification provoked by inactivation at 10 weeks of age is temporary, and resolves with a short period of binocular vision. Scale bars = 1 mm (A and D) and 50 μm (B and E). Red and blue data points indicate measurements from A and A1 layers, respectively. The timing of MI is shown in the lower left of A and D.

From the same group of animals that received binocular vision following retinal inactivation, we measured VEPs from the primary visual cortex. Consistent with our anatomical results, there was a complete recovery of visual responses after the period of inactivation wore off and animals were provided binocular vision. **Figure 7A-C** displays data from the left hemisphere of an example animal in which balanced visual responses were measured between the eyes before the right eye was inactivated for 10 days (**Figure 7A**). On the tenth day of inactivation, measurements revealed that VEPs (blue trace) were obliterated in the inactivated eye, and responses from this eye were reduced to baseline (red trace; **Figure 7B**). VEPs elicited through the formerly inactivated eye were restored to normal when 10 days of binocular vison was provided after the period of inactivation, and these responses were in balance with VEPs elicited from the other eye (**Figure 7C**).

**Figure 7.**
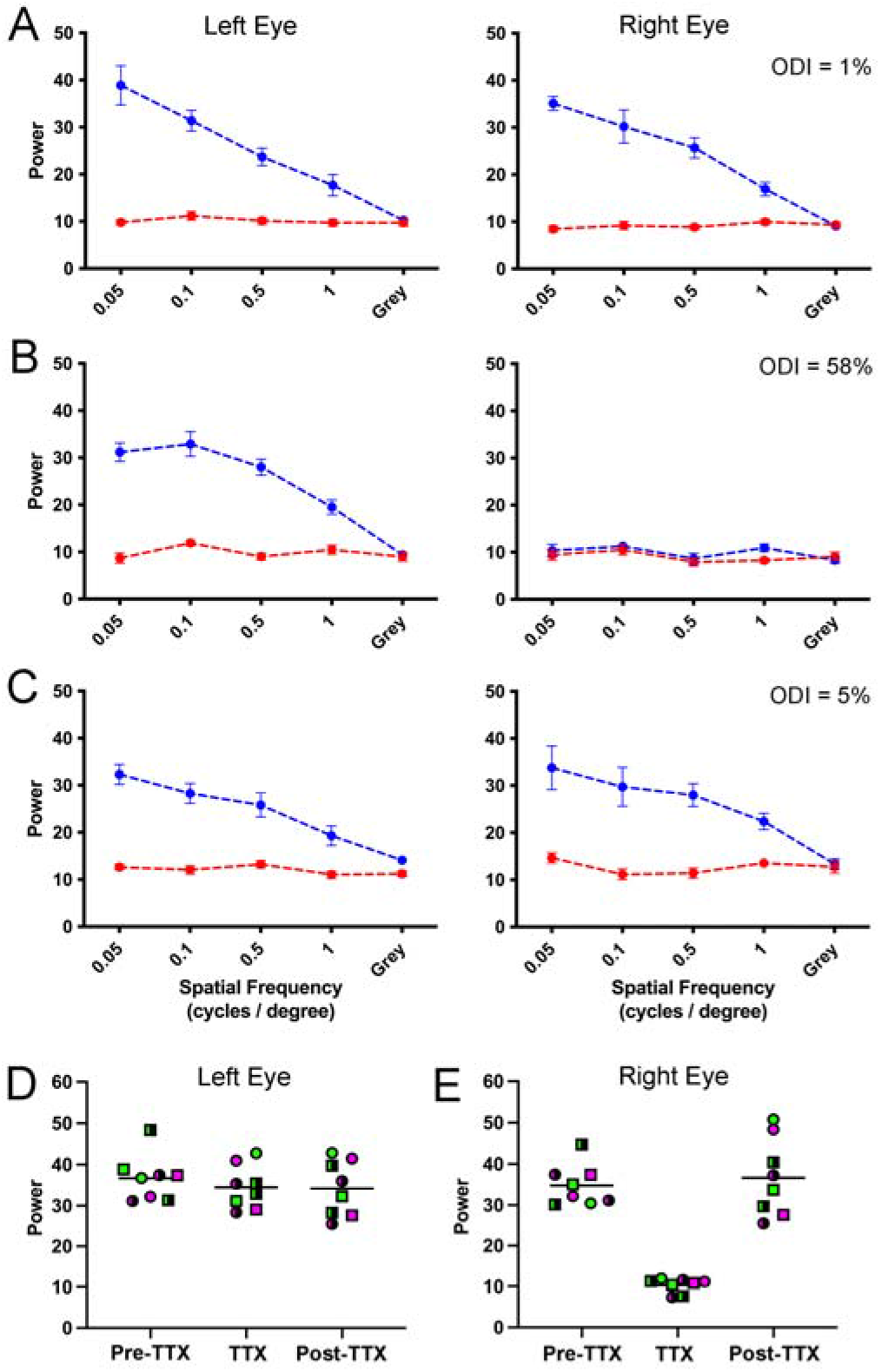
Measurement of visually-evoked potentials elicited by separate stimulation of the left and right eye before, during, and after 10 days of right eye inactivation. For each graph in A-C, spatial frequency is plotted on the abscissa, and the summed power from the Fourier analysis is plotted on the ordinate. The blue trace represents the sum of visually-evoked power, while the red trace shows the non-visual baseline power. Visually-evoked power elicited by a blank screen served as a control, and should be about equal for the blue and red trace. Data are shown for an example animal in which VEP power from the left and right eye are balanced (blue traces) prior to any visual manipulation (A). VEPs measured after 10 days of right-eye inactivation were reduced to baseline levels for that eye, while responses elicited from the left eye remained high (B). VEPs from the same animal are shown after it was provided 10 days of binocular vision following the period of inactivation (C). Restoration of normal-appearing VEPs were measured for the previously inactivated eye after the period of binocular vision, and this was in balance with VEPs measured from the left eye. In A-C, the balance of VEPs measured between the eyes is represented by an ocular dominance index (ODI; upper right value of each graph) that indicates the percentage difference between left and right eye. VEP power measurements were plotted for the left (D) and right (E) eye before (Pre-TTX), during (TTX), and after (Post-TTX) right eye inactivation for 10 days. While left eye VEP power remained unchanged across conditions, VEP power for the right eye was significantly reduced during the period of inactivation but recovered to normal levels with provision of binocular vision. Squares and circles represent data from separate animals; green and magenta represent left and right visual cortex, respectively; solid and half symbols represent data from 0.05 Hz and 0.1 Hz grating stimuli, respectively.

We next plotted VEP power measurements separately for the left (**Figure 7D**) and right (**Figure 7E**) eye for both animals before, during, and after inactivation of the right eye. In these graphs, the left and right visual cortex are represented by symbol color, different animals are represented by symbol shapes, and grating spatial frequencies are represented by symbol composition. VEP power that was measured from the normal left eye before inactivation was unchanged during or after the period of inactivation ((F(3,8) = [0.805], p = 0.418); **Figure 7D**). Inactivation of the right eye significantly reduced VEP power relative to pre-inactivation levels ((F(3,8) = [42.22], p < 0.001; multiple comparison post-hoc test: p < 0.001). The reduction of VEP power elicited through the inactivated eye recovered following 10 days of binocular vision, and was not different from VEP power measured before inactivation was imposed (multiple comparisons post-hoc test: p = 0.639). It is worth pointing out that the restoration of normal dLGN soma size and the recovery of VEPs after inactivation occurred at an age (i.e. 10 weeks old) when inactivation of the fellow eye can promote full recovery from the anatomical and physiological effects of long-term MD (Duffy et al., 2018; Fong et al., 2021). The current findings indicate that the modifications elicited by inactivation do not to have a lasting negative impact on the visual system, and instead appear to elicit a constellation of modifications that avail an opportunity for recovery from the neural modifications provoked by amblyogenic rearing.

## Discussion

In the first part of this study we compared the effect of monocular lid closure to monocular retinal inactivation as a means of assessing their respective capacities to provoke a change in soma size in the dLGN across development. Retinal inactivation for 10 days consistently produced a larger effect compared to lid closure, and this was observed at all the ages we examined. Even at ages beyond the critical period peak when MD produces little to no effect in the dLGN, MI modified soma size postsynaptic to silenced retinal ganglion cells. In addition, at all ages examined, inactivation led to cell shrinkage within both the binocular *and* monocular segments. This observation agrees with findings from Kuppermann and Kasamatsu (1983) who obtained similar results in cats after 1 week of inactivation imposed at 7 weeks of age. Thus, inactivation is qualitatively different from MD because the latter has a negligible effect on the size of neurons within the monocular segment (Guillery and Stelzner, 1970; Hickey et al., 1977). This is noteworthy because it indicates that the mechanisms inducing soma size changes between these two forms of deprivation are not identical. Moreover, despite the magnitude and breadth of the acute effects of MI, we found that the impact of inactivation can be temporary. This was determined after examining 10-week-old animals that were provided a period of binocular vision after 10 days of inactivation. At this age, the effects of retinal inactivation in the dLGN and V1 were erased a short time after the influence of TTX wore off.

Monocular retinal inactivation with TTX is known to produce a variety of neuroanatomical changes within the primary visual pathway of adult cats and monkeys. In monkey V1, loss of microtubule-associated protein occurs within ocular dominance columns that associate with the inactivated eye (Hendry and Bhandari, 1992). Loss of gamma ammino-butyric acid (GABA) and its synthesizing enzyme, glutamic acid decarboxylase (GAD), is evident in inactivated-eye ocular dominance columns of monkey V1 (Hendry and Jones, 1988), as well as within layers of the dLGN that serve the inactivated eye (Hendry and Miller, 1996). In both cats and monkeys, oxidative metabolism revealed by cytochrome oxidase staining is reduced within inactivated-eye layers of the dLGN and ocular dominance columns that serve the inactivated eye (Wong-Riley and Riley, 1983; Wong-Riley and Carroll, 1984), reflecting a decrease in neural activity that has been measured within inactivated-eye layers (Rittenhouse et al., 1999). The loss of cytochrome oxidase staining can occur quickly, being evident even after as little as 1 day of inactivation, and becomes more pronounced as the period of inactivation is extended (Wong-Riley and Carroll, 1984). Findings from the current study, in agreement with those from Kuppermann and Kasamatsu (1983), demonstrate that MI significantly reduces dLGN cell size within layers connected to the affected eye. The effect of MI is evident within both the binocular and monocular segments, which distinguishes it from MD because the latter produces alterations only within the binocular segment. Therefore, the effect of MI is unlikely to be explained by a competitive imbalance between the eyes because it is observed within the monocular segment where geniculostriate axons serving the inactivated eye are not subjected to competition with those from the fellow eye. This raises the possibility that the effects of MI on the dLGN are catalyzed directly by an antecedent loss of tonic retinal ganglion cell activity.

A possible contributor to the postsynaptic somatic changes induced by inactivation may be modification of trophic support provided by the retina. Trophic support of dLGN neurons is contributed by selective anterograde transport of neurotrophins from the retina in an activity-dependent manner (Caleo et al., 2000). The anterograde transport and release of brain-derived neurotrophic factor (BDNF) from retinal ganglion cells can elicit expression of immediate early genes (c-fos and Zif268) within target neurons of the dLGN (Caleo et al., 2000). In cats, MI for 2 days with intravitreal microinjection of TTX significantly lowers BDNF levels within layers of dLGN, as well as in V1 ocular dominance columns serving the inactivated eye (Lein and Shatz, 2000). Although it is widely thought that deprivation-induced ocular dominance plasticity is cortically mediated, augmenting levels of retinal BDNF depleted by visual deprivation can mitigate the typical cortical effects of MD (Mandolesi et al., 2003), indicating a possible role for the retina to influence neural plasticity throughout the primary visual pathway. It therefore seems possible that the shrinkage of cells in the dLGN following MI may involve a reduction of trophic support from the retina, which would presumably affect the binocular and monocular segments equally. The effect of inactivation observed in the current study was clearly reduced with age, which implies that the dependence of dLGN neurons on activity-dependent trophic support is likewise regulated by age. Irrespective of the mechanisms involved, that inactivation retained potency to evoke a change in cell size at older ages than is possible with MD suggests that it adheres to a longer-lasting critical period that could be leveraged to prolong the reversibility of neural dysfunctions in the primary visual pathway.

Monocular retinal inactivation in the youngest animals we examined produced an effect in the dLGN binocular segment that was about double the magnitude of what was measured in the monocular segment (**Figure 5**). However, at older ages the size of the inactivation effect in the binocular segment was about the same as that measured in the monocular segment. In accordance with Sherman and Spear (1982), a combination of binocular competition and noncompetitive mechanisms is proposed when neural abnormalities occur within both dLGN segments but are more severe in the binocular segment. When abnormalities are equally severe in both segments, a noncompetitive mechanism is suggested. Therefore, the larger inactivation effect we observed in the dLGN binocular segment at younger ages near the critical period peak may originate from a combined influence of cortically-mediated binocular competition between eye inputs, in conjunction with noncompetitive mechanisms that may include a modification of trophic influence from the retina. The comparable effect magnitude that was measured between the binocular and monocular segments when inactivation occurred at older ages suggests that the effect of inactivation later in development derives from noncompetitive mechanisms that might reasonably be attributed to loss of tonic retinal activity.

When MI occurred at 10 weeks of age, its impact on dLGN cell size was significant and comparable to that elicited by an equal duration of MD imposed at the peak of the critical period (Kutcher and Duffy, 2007). Notwithstanding this large effect, subsequent provision of a short duration of binocular vision was sufficient to reverse the atrophy of cells. Likewise, inactivated-eye VEPS that were reduced to baseline during the period of inactivation, returned to normal a short duration after the final TTX injection. Even when MI occurs at 7 weeks of age, a time when plasticity capacity is quite high, the effect of MI on dLGN soma size recovers with subsequent binocular vision (Kuppermann and Kasamatsu, 1983). Within the primary visual pathway of cats and monkeys subjected to MI, the significant reductions of cytochrome oxidase staining (Wong-Riley and Riley, 1983; Hendry and Miller, 1996) as well as GAD and GABA immunolabeling (Hendry and Jones, 1988) all recover and are indistinguishable from normal when inactivation is followed by a period of binocular vision. This indicates that inactivation does not set into motion an irreversible degenerative process that would cause permanent alteration, which distinguishes it from monocular enucleation that does elicit degeneration and loss of neurons in the dLGN (Kalil, 1980). Although we did not directly examine whether the spontaneous recovery measured after MI at 10 weeks of age also occurs after an equal period of MD that is relieved at the same age, a prior study from our group provides some insight. We previously documented incomplete dLGN recovery when 10 days of binocular vision followed 6 weeks of MD started at 4 weeks of age (Duffy et al., 2018). In other words, while the effect of MD did not resolve when binocular vision was provided at 10 weeks of age, the effect of MI did. Although in this comparison the timing of MD was different than that of MI, it raises the possibility of an additional distinction between MI and MD, namely, that the effects of MI are temporary and can resolve spontaneously with the restoration of binocular vision. It is important to point out that although the transient effects of inactivation on characteristics of neurons within dLGN and V1 have been documented in cats as young as 7 weeks old, the reversible nature of these alterations may not be observed when applied at much younger ages. Indeed, abnormal receptive field properties of kitten dLGN neurons has been documented following 5-8 weeks of inactivation when started neonatally (Archer et al., 1982), suggesting that abolition of retinal output can promote development of abnormal retino-geniculate connections when it occurs near the time of birth.

Following a long-term MD in which ocular dominance is shifted strongly in favor of the non-deprived eye, inactivation of the stronger eye in both mice and cats can promote recovery of visually-driven cortical responses elicited by the weaker eye (Fong et al., 2021). Similar recovery occurs in the cat dLGN, where deprived neurons rendered smaller by MD recover to their normal size when the non-deprived eye is inactivated for 10 days (Duffy et al., 2018). This recovery occurs without any detectable systemic toxicity, and several studies that have administered intravitreal TTX in cats and monkeys have reported no evidence of ocular histopathology or disruption to the eye’s normal function once the period of inactivation has passed (Wong-Riley and Riley, 1983; Wong-Riley and Carroll, 1984; Foeller and Tychsen, 2019; Dicostanzo et al., 2020). Consistent with results from the current study showing that MI can elicit a significant modification of cell size at older ages than is possible with MD, recovery from the effects of a long-term MD using fellow-eye inactivation can occur at an age when the same duration of reverse occlusion fails to produce significant recovery (Duffy et al., 2018). Given our current findings that indicate the effect of MI at 10 weeks of age appears to originate from noncompetitive mechanisms, it possible that the inactivation-induced recovery from long-term MD is also due to noncompetitive mechanisms, rather than by creating a competitive imbalance between the eyes that occurs with occlusion therapy. It remains to be determined if the morphological effects of MI are obligatory for the functional recovery from MD. Nevertheless, the results from the current study have delineated potentially important characteristics of inactivation that may contribute to its unique potency to promote neural plasticity. It will be important to investigate the capacity for inactivation to promote recovery from MD at even older ages than have been examined to date.

## Notes

### Competing Interest Statement

The authors have declared no competing interest.

